# Statistical significance of cluster membership for determination of cell identities in single cell genomics

**DOI:** 10.1101/248633

**Authors:** Neo Christopher Chung

## Abstract

Single cell RNA sequencing (scRNA-seq) allows us to dissect transcriptional heterogeneity arising from cellular types, spatio-temporal contexts, and environmental stimuli. Cell identities of samples derived from heterogeneous subpopulations are routinely determined by clustering of scRNA-seq data. Computational cell identities are then used in downstream analysis, feature selection, and visualization. However, how can we examine if cell identities are accurately inferred? To this end, we introduce non-parametric methods to evaluate cell identities by testing cluster memberships of single cell samples in an unsupervised manner. We propose posterior inclusion probabilities for cluster memberships to select and visualize samples relevant to subpopulations. Beyond simulation studies, we examined two scRNA-seq data - a mixture of Jurkat and 293T cells and a large family of peripheral blood mononuclear cells. We demonstrated probabilistic feature selection and improved t-SNE visualization. By learning uncertainty in clustering, the proposed methods enable rigorous testing of cell identities in scRNA-seq.

## Introduction

Recent high-throughput genomic technologies, such as single cell RNA-seq (scRNA-seq), have increased a number of single cell samples that can be profiled simultaneously (Navin et al., 2011; Xu et al., 2012; Shalek et al., 2013; Wills et al., 2013). With a large number of gene expression profiles of single cells, we are often interested in their cellular heterogeneity. Encompassing cellular types, spatio-temporal contexts, and environmental stimuli, cellular heterogeneity can be characterized from systematic variation in gene expression profiles. Routinely, unsupervised clustering of scRNA-seq data approximates *K* subpopulations among *m* single cell samples, determining individual cell identities. Computationally defined *m* cell identities are then used in downstream data analysis and exploration. Given that *m* cell identities in *K* subpopulations are determined by unsupervised clustering, it is critical to test if individual *m* cell identities are correctly assigned. We have developed novel methods to estimate statistical significance and posterior probabilities of assigning cell identities to such subpopulations. Using simulated and scRNA-seq data, the proposed statistical tests for cluster membership are demonstrated to identify relevant samples belonging to subpopulations, that provide intuitive improvements in feature selection and visualization.

Clustering has been one of the most popular analysis methods for high-dimensional genomic data. Microarray studies pioneered use of clustering to identify co-regulated subsets of genes (Spellman et al., 1998; Eisen et al., 1998; Gasch et al., 2000) and subpopulations among samples (Alon et al., 1999; Golub et al., 1999; Sørlie et al., 2001). In absence of precise labels about samples, *m* gene expression profiles could be clustered into *K* subpopulations with differential responses in an unsupervised manner (Alon et al., 1999; Golub et al., 1999; Sørlie et al., 2001). Recently, there have been several scRNA-seq studies where genome-wide gene expression from thousands of single cells are measured en masse (Patel et al., 2014; Jaitin et al., 2014; Macosko et al., 2015; Zheng et al., 2017). Unlike microarray or bulk RNA-seq data that may average gene expression from a large number of cells, cellular heterogeneity is now observed at a single cell resolution. Such heterogeneity is characterized by clustering *m* single cell samples to *K* subpopulations, providing computationally defined *m* cell identities (Figure 1). These cell identities, which may be related to tissue types, cell lineage/cycle, and other molecular processes, are widely used in scRNA-seq analysis. To this end, a number of clustering algorithms have been applied and developed.

**Figure 1:**
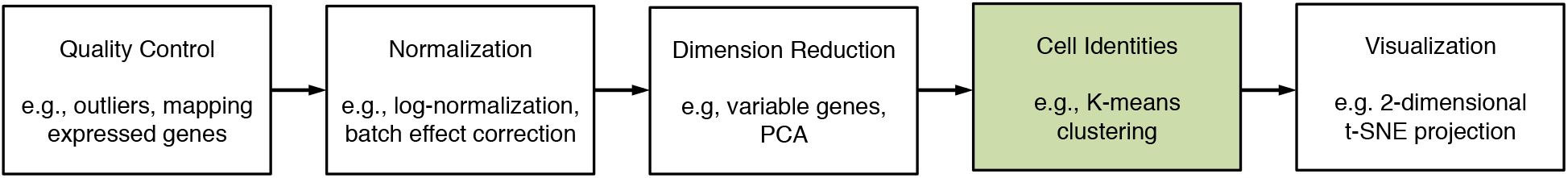
Illustration of scRNA-seq analysis steps. Unsupervised clustering is often used to computationally determine cell identities. The proposed methods enable statistically rigorous evaluation of computational cell identities.

Single cell analysis tools implement various clustering algorithms, including, but not limited to, K-means clustering in Seurat (Satija et al., 2015; Butler et al., 2018), hierarchical clustering in SINCERA (Guo et al., 2015), and density peak clustering in Monocle (Rodriguez and Laio, 2014; Qiu et al., 2017). Furthermore, a number of clustering methods specifically tailored to scRNA-seq data have been developed to identify subtypes of single cells (Zeisel et al., 2015; Xu and Su, 2015; Buettner et al., 2015; Wang et al., 2017). To increase computational efficiency and signal-to-noise ratios, a number of scRNA-seq studies combine principal component analysis (PCA; Jolliffe (2002)) or t-Distributed Stochastic Neighbor Embedding (t-SNE; van der Maaten and Hinton (2008)) with unsupervised clustering (Macosko et al., 2015; Satija et al., 2015; Zheng et al., 2017). Complementarily, there are consensus (ensemble) algorithms that combine different clustering results such as SC3 that utilizes multiple results of K-means clustering with different transformations (Kiselev et al., 2017). Due to its computational efficiency and general purpose, K-means clustering remains exceptionally popular in scRNA-seq algorithms and applications. Therefore, we focus on K-means clustering and its variants - Partitioning Among Mediods (Kaufman and Rousseeuw, 1987), mini batch K-means (Sculley, 2010), and K-means++ initialization (Arthur and Vassilvitskii, 2007) for the proposed statistical test of cluster membership. Nonetheless, the proposed methods may be readily incorporated into other clustering algorithms and single cell analysis tools.

After clustering samples into *K* clusters representing *K* subpopulations, we are interested in evaluating reliability of computational cell identities and subsequently identifying significant single cell samples belonging to each subpopulation. Our innovative data resampling and testing scheme for unsupervised classification evaluate whether observed samples are true members of corresponding clusters. Complementing the jackstraw methods for PCA and related methods (Chung and Storey, 2015), we automatically learn and account for overfitting inherent in using cluster structure estimated from scRNA-seq data. By estimating p-values and posterior inclusion probabilities (PIPs), the proposed jackstraw tests help identify and visualize samples that are the most relevant for data-dependent subpopulations. This connects unsupervised classification of high-dimensional data and fundamental hypothesis framework in a statistically rigorous, yet straightforward, manner. Our non-parametric approach does not require modifications to clustering algorithms or transformations of scRNA-seq data.

We used simulated and scRNA-seq data to demonstrate operating characteristics of the proposed methods. In the first part of simulation studies, a series of datasets were generated from a latent variable model, while increasing an amount of noise. The joint behavior of p-values are scrutinized by conducting 100 independent simulations. In the second part of simulation studies, we used a cluster structure from scRNA-seq data of 2638 peripheral blood mononuclear cells (PBMCs) by 10X Genomics Chrominum. Two scRNA-seq applications are presented using a mixture of Jurkat and 293T Cell Lines and 68579 PBMCs (Zheng et al., 2017). We further implemented a scalable mini batch version of K-means clustering for tens of thousands samples. By applying the proposed methods on unlabeled single cell samples, we show improved classification and visualization of cell identities. These proposed methods are implemented in a R package called *jackstraw* (Stable Release: https://CRAN.R-project.org/package=jackstraw and Development: https://github.com/ncchung/jackstraw).

## Methods and Algorithms

The observed data **Y**_(*m,n*)_ contains *m* rows and *n* columns. In scRNA-seq data, we assume that single cell samples are arranged as rows, whereas columns as genes. While scRNA-seq generates counts of sequence reads, a variety of tools (Satija et al., 2015; Guo et al., 2015; Qiu et al., 2017; McCarthy et al., 2017) are used for quality controls and normalization (Figure 1). Furthermore, feature selection and dimension reduction may be applied to highlight certain aspects of systematic variations or biomarkers of interest. For simplicity, genomic variables in *n* columns are interchangeable denoted as genes. Lastly, even though we focus on clustering single cell samples in rows, it is trivial to apply on columns as well.

Consider there exists *K* clusters (i.e., subpopulations) where *m_k_* samples are assigned to *k*^th^ cluster (*k* = 1,…, *K*). Then, *m*_1_,…,*m_K_* samples are assigned into corresponding 1,…,*K* clusters, respectively, where 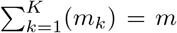. Samples within *k*^th^ cluster exhibit systematic variation that may be summarized by their center **c**_*k*_ for *k* = 1, … *K*. For example, in K-means clustering, the mean is the center where the nearest center in Euclidean (*L*_2_) distance are used to assign observed samples to the clusters. In Partitioning Among Mediods (PAM), the center is always selected from observed samples and *L*_1_ distance is used for membership assignments (Kaufman and Rousseeuw, 1987). If **y**_*i*_ is assigned to *k*^th^ cluster with **c**_*k*_, its membership indicator *β_i,k_* is 1. By definition, the subset of samples **y**_*i*_ with *β_i,k_* = 1 make up **c**_*k*_.

Cluster structure may be viewed as distinct patterns being manifested on subpopulations of samples. Among single cells, clusters are naturally presumed to reflect cell identities. Cell identities may arise from cellular processes whose distinct systematic characteristics are embedded in gene expression of those subpopulations. In a subpopulation that are expected to be clustered together, there are latent variables l_*k*_ and dichotomous membership indicators **b**_*k*_ for *k* = 1,…,*K*. **L** may assume a wide range of patterns including continuous or categorical structures (Linda M. Collins, 2010; Bartholomew et al., 2011). Coupled with independently and identically distributed noise **E**, the data may be modeled as:

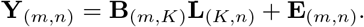

If a particular *i*^th^ sample is truly associated with a **l**_*k*_, its coefficient *b_i,k_* is 1. Otherwise, 0. Row-wise means may be handled by centering the data, whereas row-wise variances preserved by the proposed resampling scheme.

We use this model for simulation studies with the null samples that are independently and identically distributed. Generally, p-values corresponding to null samples should be uniformly distributed between 0 and 1. The critical operating characteristics of a statistical test is to maintain this uniformity, which can be tested in simulation by a Kolmogorov-Smirnov (KS) test. To reflect complexity of scRNA-seq data, we also consider cluster structure directly extracted from gene expression profiles of PBMCs (Download available from https://support.10xgenomics.com/single-cell-gene-expression/datasets).

There have been important developments in unsupervised learning that consider mixture or latent variable models that improve our understanding and interpretation of data (Yeung et al., 2001; McLachlan and Peel, 2004; Fraley and Raftery, 2007). However, even model-based clustering approaches or regularization do not provide cluster centers and membership assignments that can be used again with the observed data, resulting in so-called “double dipping.” Our approach learns and incorporates inevitable uncertainty in assigning single cell samples to clusters, that are directly derived from the same set of samples. Unlike fuzzy or probabilistic clustering algorithms, the proposed statistical tests enhance unsupervised clustering without requiring modifications to well-established clustering algorithms.

### Jackstraw Data

We illustrate the proposed approach to construct and utilize a minimally disruptive data that maintain the overall subpopulations. At the same time, the jackstraw data include synthetic null samples that truly do not belong to any subpopulation. Operationally, out of *m* observed samples, a relatively small number (*s* ≪ *m*) of samples are resampled without replacement, resulting in s synthetic null samples. Other *m* − *s* observed samples are unchanged. The jackstraw data **Y**\* combines s synthetic null samples and intact *m* − *s* observed samples (Figure 2). Subpopulations are largely preserved in the jackstraw data, because a relatively small number of observed samples became i.i.d. noise.

**Figure 2:**
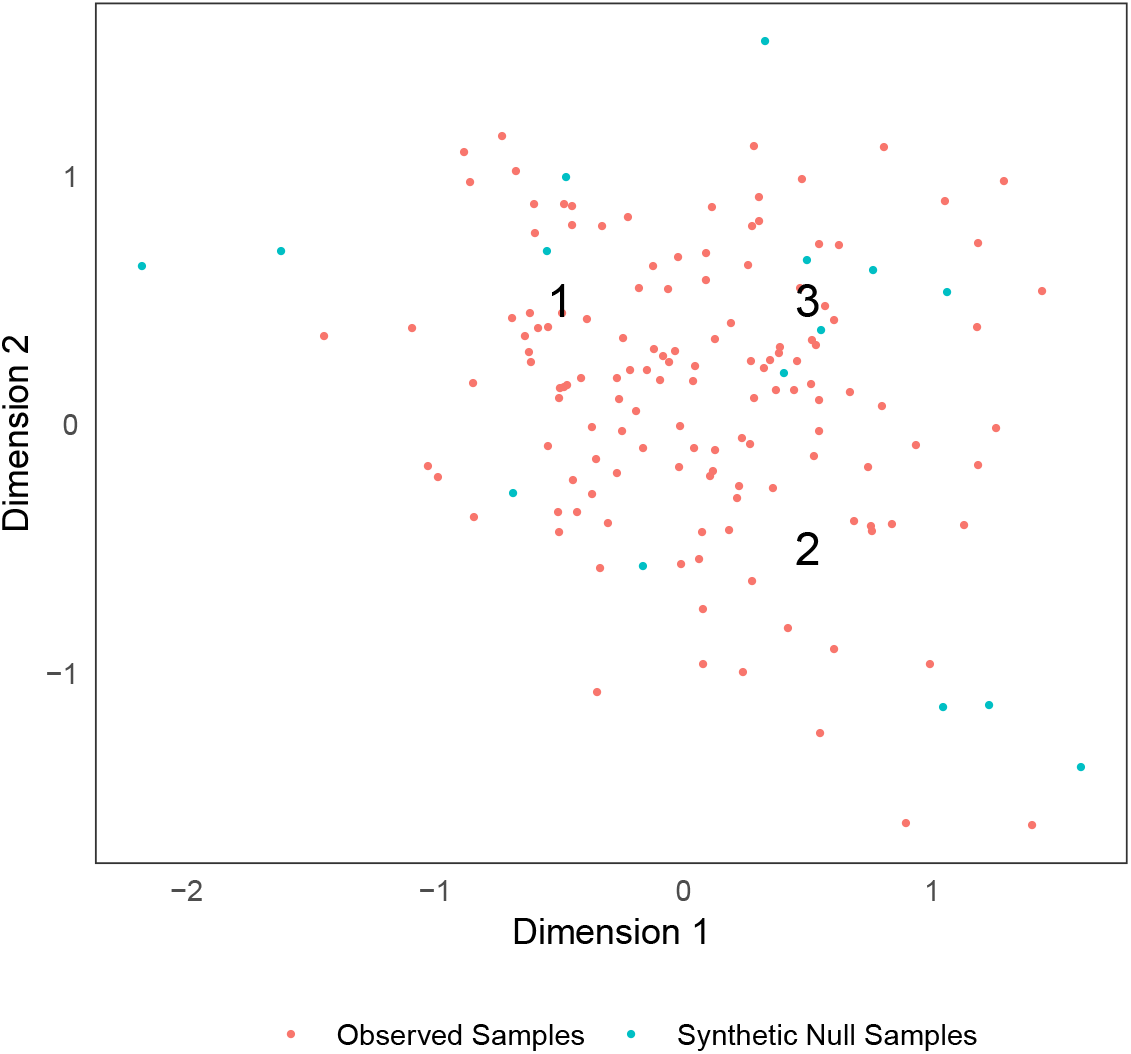
Illustration of the jackstraw data **Y**\* consisted of *s* synthetic null samples and *m* − *s* observed samples. Among 135 observed data points shown, there exist *K* = 3 clusters, denoted by numbers at their centers. Fifteen blue points are synthetic null samples that are i.i.d. The proposed methods leverage this information to learn overfitting characteristics in clustering.

Falling in the broad family of resampling strategies, this jackstraw procedure is distinct from the jackknife (Quenouille, 1949; Tukey, 1958) and the bootstrap (Efron, 1979). Developed to non-parametrically estimate variance and bias, the jackknife would estimate a statistics on *m* − 1 samples after randomly removing one sample. The bootstrap would resample *m* samples with replacement. The bootstrap and similar ideas has been used to evaluate stability or prediction strength in unsupervised clustering (Jain and Moreau, 1987; Kerr and Churchill, 2001; Ben-Hur et al., 2002; Tibshirani and Walther, 2005; Hennig, 2007). These methods, that have often been developed in microarray studies, are interested in evaluating clusters themselves. Furthermore, in cross-validation where *m* samples are split into training and test sets, one may cluster a training set to obtain cluster centers and use a test set to compute distances. However, as hold-out samples are not used to estimate cluster structure, this can not learn or account for overfitting.

In contrast, we aim to minimally disrupt the observed data, such that a few synthetic null samples are both used for clustering and estimating the empirical null distribution. When we cluster the jackstraw data, cluster centers 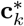 of **Y**\* are almost identical to the original cluster centers **c**_*k*_ of **Y** (for *k* = 1,…, *K*). Because of the nature of clustering algorithms, all samples in **Y**\*, including *s* synthetic null samples, will be assigned to one of *K* clusters. When a synthetic null sample 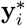 is assigned to *k*^th^ cluster, an association statistics between 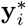 and 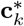 is under the null model that assumes independence since 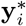 is i.i.d. by definition. Yet, because 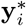 is stochastically assigned to 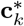, we effectively learn the overfitting characteristics of clustering. Over a large number of iterations *b* =1, …,*B*, the empirical distribution of null statistics is formed. This empirical distribution of null statistics is used to evaluate significance of individual samples.

Evaluation of cluster membership requires a pre-defined number of clusters *K*. There is a vast amount of literature on the choice of *K*, which is beyond the scope of this study. Methods, including aforementioned bootstrap procedures, have been proposed in the last five decades in this area of research including cluster stability and reliability statistics (Akaike, 1974; Schwarz et al., 1978; Bock, 1985; Fraley and Raftery, 1998; Pelleg et al., 2000; Tibshirani et al., 2001; Hamerly and Elkan, 2004; Liu et al., 2008; Huang et al., 2015). Exploration of the observed data requires understanding of experimental designs, prior biological (domain) knowledge, and visual analytics.

### Jackstraw Tests for Cluster Membership

K-means clustering is one of the most established algorithms (MacQueen et al., 1967; Hartigan and Wong, 1979; Lloyd, 1982). K-means clustering is the most efficient algorithms (Wong, 2015). Particularly, considering rapidly growing sizes of scRNA-seq data, hierarchical clustering, graph-based community detection, and density-based clustering are shown to be much slower than K-means clustering (Fan, 2018). Due to its efficiency and well-known characteristics, it has been popularly used in a number of scRNA-seq studies to computationally determine cell identities (Macosko et al., 2015; Satija et al., 2015; Zheng et al., 2017). Furthermore, we implement Partitioning Around Medoids (PAM) (Kaufman and Rousseeuw, 1987) and mini batch K-means clustering (Sculley, 2010; Sanderson and Curtin, 2016) for robustness and scalability. Lastly, when possible, we use a K-means++ initialization algorithm to speed up our clustering steps (Arthur and Vassilvitskii, 2007).

We present a detailed jackstraw method to estimate statistical significance of cluster memberships when using K-means clustering and their variants. In the algorithm 1, we use F-statistics where the full models include appropriate cluster centers. The use of F-statistics allows us to flexibly specify the full and null models, which may incorporate other covariates in more complex settings. However, depending on distributional assumptions or computational needs, one may devise to use other association statistics.

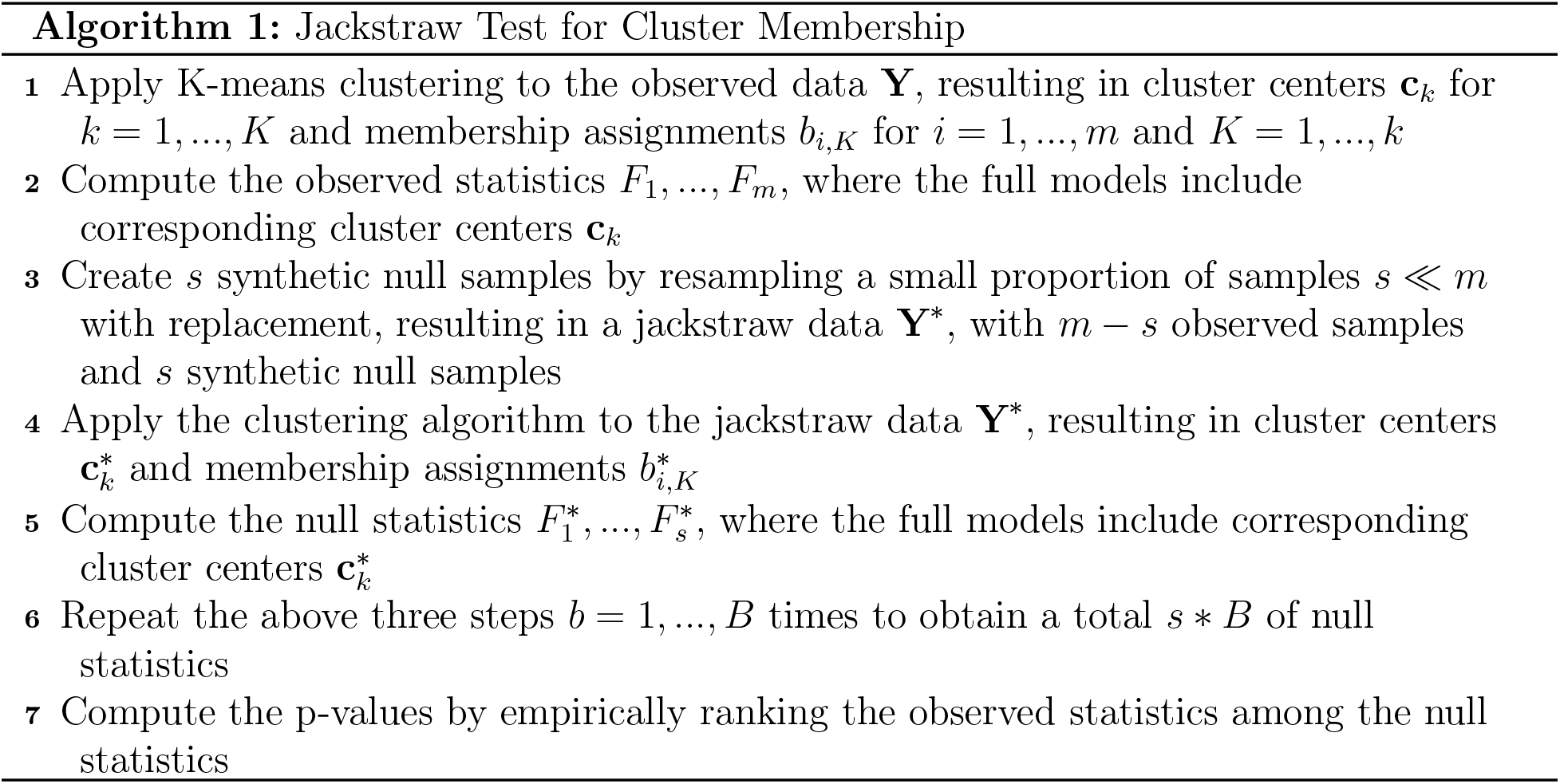

The choices of *s* and *B* control the speed of computation, while the total number of null statistics (*s* × *B*) determines the overall p-value resolution. For *B* iterations, we need to cluster the jackstraw data *B* times, and for each iteration *b* =1,…, *B*, we obtain *s* null statistics. Assuming *s* × *B* is hold constant, a smaller *s* provides more accurate p-values, while increasing computational burdens. Therefore, we want to ensure the original clusters are preserved as much as possible, permitting the computational power. As we increase the number of synthetic null samples *s* in **Y**\*, the overall systematic variation captured by *K* cluster centers may be increasingly disrupted. While we use *s* ~ 0.1 × *m* for genomic data, the number of clusters (*K*) and the proportion of samples assigned to them (*m*_1_,…, *m_k_*) must be considered. A higher value of *K* for a given *m* would need a smaller *s*, so that the clusters with limited members are represented in the jackstraw data.

The overwhelming disruption would further inflate null F-statistics, since a larger number of synthetic null samples would make up 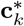. In extreme scenarios where all samples have been resampled, the new cluster centers are completely dominated by independent synthetic null samples. This operating characteristic allows us to guard against artificially inflated significance and to guide the input parameters for the proposed algorithm. In practice, we input **C** as the initial centers for *K* clusters when clustering the jackstraw data for efficient convergence.

In contrast, the conventional resampling methods can be applied to the cluster centers, resulting in a “naive” significance test. After all *m* samples are resampled with replacement, their F-statistics with respect to **c**_*k*_ are used to form an empirical distribution of null statistics. Observed F-statistics are compared to this empirical distribution to obtain naive p-values. This circular analysis inflates statistical significance, since the observed samples are used twice to compute the cluster centers and to again test against the cluster centers. Essentially, this represents how the bootstrap or the permutation would be applied to cluster membership assignments. We apply the conventional methods in simulation studies to demonstrate how the jackstraw approach overcomes this type of overfitting.

Tens of thousands of single cell samples may prove to be too burdensome for K-means clustering. Therefore, we incorporate a scalable mini batch version of K-means (Sculley, 2010; Sanderson and Curtin, 2016) using `ClusterR` package, where a random subset of data are used iteratively to update cluster centers and membership assignments (Step 1 and 4 in the algorithm 1). Similarly, instead of randomly selecting cluster centers, K-means++ initialization may improve its convergence, which is available as an option for our implementation (Arthur and Vassilvitskii, 2007). Because K-means clustering relies on Euclidean distance, one may be concerned about its robustness to outliers or generalizability to other distributions. By choosing observed data as cluster centers and using *L*_1_ norm, Partitioning Around Medoids (PAM) may perform more appropriately and include in our implementations (Kaufman and Rousseeuw, 1987) using cluster package.

### Posterior Inclusion Probabilities

After clustering *m* single cell samples into *K* subpopulations, the proposed jackstraw method estimates a probability that an individual sample may have been assigned to a given subpopulation by chance. For intuitive feature selection and visualization, we investigated how to harness *m* p-values (or, the empirical null distribution) to filter, de-noise, and visualize the clusters. When considering high-dimensional samples typical in scRNA-seq studies, it is advantageous to consider a family of multiple hypotheses simultaneously (Efron, 2012). Particularly, from jackstraw p-values, we propose to calculate posterior probabilities that samples are truly included in a given cluster. A discussion of posterior inclusion probabilities (PIPs) that are used for shrinkage and improvement of latent variable estimates is available in *Chapter 3* of (Chung, 2014).

Consider that the *m* jackstraw p-values **p** = *p*_1_,…,*p_m_* are obtained for *m* single cell samples that have been clustered into *K* subpopulations. We are interested in estimating a posterior probability that *b_i_* ≠ 0, since non-zero coefficients imply their bona fide inclusion in the cluster:

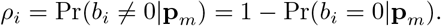

PIP can be readily obtained by estimating Pr(*b_i_* = 0|**p**_*m*_) through an empirical Bayes approach (Efron et al., 2001; Efron, 2007). In multiple hypothesis testing, Pr(*b_i_* = 0|**p**_*m*_) is called a local false discovery rate (FDR). With a large amount of samples, it may be advantageous to consider posterior probabilities among each subpopulation or to improve estimation of the proportion of null samples (*π*_0_) using prior biological knowledge. There also exist related Bayesian methods that could be explored for specific applications and prior knowledge (Barbieri and Berger, 2004; Scott and Berger, 2005; Ghosh et al., 2006).

These results in *m* PIPs for *K* subpopulations, that can be used for:

1. Retaining a subset of samples **y**_*i*_ with ***ρ***_*i*_ > *α_ρ_*, where *α_ρ_* is a user-defined threshold,
2. Visualizing samples in reduced dimensions (e.g., PCA, t-SNE) where transparency ~ ***ρ***_*i*_,
3. Improving the cluster centers by weighting the corresponding samples with ***ρ***_*i*_.

Local FDRs and PIPs from *K* subpopulations of multiple hypothesis tests can be flexibly combined for downstream analyses, as to aid feature selection and dimension reduction. When applying the proposed methods on computational determination of cell identities in scRNA-seq data, we incorporate PIPs to hard-threshold and soft-threshold the observed single cell samples. After having obtained PIPs, samples with low PIPs can be removed (i.e., hard-thresholding), achieving feature selection. A subset of samples above a certain PIP threshold (e.g., PIP > 0.8) may be visualized in 2-dimensional PCA or t-SNE projections. Alternatively, one may use PIPs to automatically control transparencies or colors that would emphasize samples with high PIPs. Furthermore, this approach may improve a wide range of clustering, such as improved assignments of single cell samples to subpopulations and regularization of cluster centers.

## Simulation Studies

To demonstrate the operating characteristics of the proposed statistical tests, we conducted two sets of simulation studies, which enabled critical assessment of p-values using null samples that are known to be i.i.d. noise. First, we generated a data from relatively simple latent variable models where we change an amount of noise 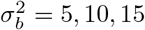. By simulating 100 independent cases and testing a joint distribution of null p-values, we showed that the proposed methods meet the joint null criterion, which is a stringent benchmark for simultaneously testing multiple hypotheses (Leek and Storey, 2011). Second, to reflect complexity of scRNA-seq data, we considered a cluster structure from gene expression profiles of 2700 peripheral blood mononuclear cells (PBMCs). Essentially, 8 clusters with varying amounts of signals and assigned samples are used to simulate according to an aforementioned model.

### Evaluation using Joint Null Criterion

In the first set of simulation studies, we generate *m* = 1000 samples (rows) and *n* = 100 genes (columns) following the latent variable model described in *Methods and Algorithms*. Latent variables L are drawn from the Normal(*μ* = 0, *σ*^2^ = 1) distribution. Relationships between **l**_*k*_ and samples are given by dichotomous coefficients **B** where *b_i,k_* indicates whether **y**_*i*_ is a member of **l**_*k*_ for *k* =1,…, *K* and *i* =1,…, *m*. The noise **B** is drawn i.i.d. from 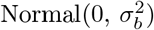, where its variance governs the noise level. Forming *Oracle Group A*, the 500 samples are true members of the signal cluster arisen from **l**_1_ with *b*_*i*,1_ = 1 for *i* = 1,…, 500. The other 500 samples are i.i.d. noise, in *Oracle Group B*, which can be viewed as being centered around the *n*-dimensional origin. Therefore, a true proportion of null smaples is *π*_0_ = .50.

We simulated three scenarios using 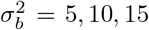 as an increasing noise level brings these two groups closer and makes the clustering task more difficult. PCA was applied on the dataset realized from each configuration to visualize the top 2 PCs (Figure S1). Without knowing or using *Oracle Groups*, the proposed jackstraw tests were applied for K-means clustering. Theoretically, the null p-values from the samples that are not related to the latent variables (corresponding to *Oracle Group B*) should form the Uniform(0,1) distribution, which can be evaluated by the Kolmogorov-Smirnov (KS) test. We repeated a given simulation configuration 100 times independently and investigated how 100 KS test p-values from 100 independent simulations meet the joint null criterion (Leek and Storey, 2011).

We describe one simulation from the main scenario involving a moderate amount of noise 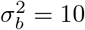. While 1000 samples were split almost equally between *Cluster 1* and *2*, 30 and 470 null samples were members of *Cluster 1* and *2*, respectively. Because *Cluster 1* contained 470 samples related to the latent variable l_1_, its center and l_1_ were highly correlated with a Pearson correlation of 0.99. The proposed test was then applied on the simulated data with *s* = 100 synthetic null samples over *B* = 5000 iterations. Figure 3(a) shows histograms of p-values stratified by *Oracle Groups* as parametrized by **B**. In *Oracle Group B*, the jackstraw p-values corresponding to 500 null samples are uniformly distributed between 0 and 1. In contrast, the naive significance tests are highly anti-conservative, pushing towards 0. In *Oracle Group A*, the jackstraw p-values are greater than the naive p-values because the jackstraw approach learns the overfitting characteristics and fixes an anti-conservative bias (Figure 3(a)). Utilizing all *m* p-values, the proportion of null samples are estimated to be 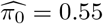 for the jackstraw and 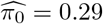 for the naive methods.

**Figure 3:**
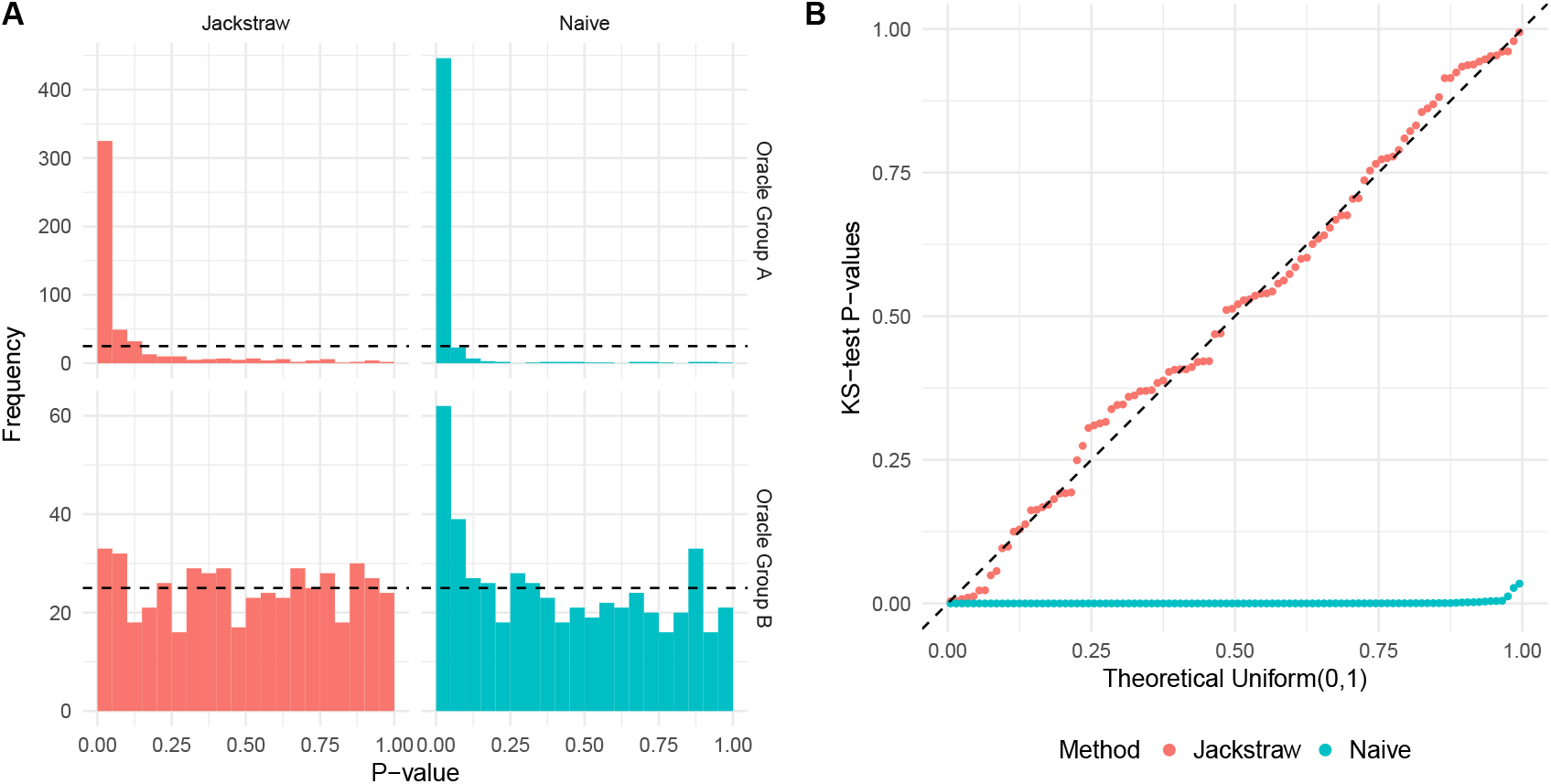
Evaluation of the jackstraw tests for clustering using the main simulation study with *σ* = 10. *Oracle Group A* contains 500 samples that are derived from a latent variable l 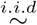 Normal(0,1), whereas 500 samples in *Oracle Group B* are noise. *(a)* Histograms of p-values stratified by methods. The jackstraw and tests were applied without using any simulation parameters. *(b)* This simulation study was repeated 100 times, where null p-values corresponding to *Oracle Group B* were evaluated against Uniform(0,1) distribution using Kolmogorov-Smirnov (KS) tests. QQ plot of KS p-values from the jackstraw and naive methods are shown, where valid p-values follow a diagonal line.

We repeated this configuration to ensure accuracy and robustness across 100 independent simulations. In each simulation, we examined the joint behavior of 500 null p-values from *Oracle Group B* using a double-sided KS test. When the joint behavior of those KS test p-values follows the i.i.d. Uniform(0,1) distribution (where the double KS test p-value > *α_jnc_*), the subsequent multiple hypothesis testing procedures, including false discovery rates, hold true (Leek and Storey, 2011). In other words, meeting the stringent standard of the joint null criterion demonstrates that the proposed methods overcome circular analysis inherent in utilizing data-dependent cluster centers and membership assignments and that the p-values are jointly and marginally accurate. One hundred KS test p-values, estimated from both the jackstraw and naive methods, are visualized against the Uniform(0,1) distribution (Figure 3(b)). The jackstraw tests satisfy the joint null criterion, where 100 KS test p-values are uniformly distributed (double KS test p-value = 0.79). In contrast, the naive methods are strongly anti-conservative, where 100 KS test p-values are strongly skewed towards 0 (double KS test p-values < 2.2 × 10^−16^).

Results from two additional simulation configurations 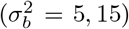, that are both repeated 100 times, are shown in Figure S2 and Figure S3.

### Simulation using Clusters of scRNA-seq Data

We used single cell gene expression of PBMCs generated by Chrominum (v1 Chemistry) to produce a complex simulation study with scRNA-seq characteristics. In particular, we obtained a dataset called `pbmc3k` from 10X Genomics^1^, which originally contains 2700 PBMC samples. Although PBMCs are composed of multiple subtypes with distinct characteristics, a Chrominum platform outputs unlabeled scRNA-seq data such that individual cell identities are unknown. The cell identities are often computationally defined from a clustering analysis of gene expression data. In our PBMC simulation study, we generate a gene expression dataset based on a cluster structure of normalized `pbmc3k` data, that is subsequently used to evaluate the jackstraw methods.

We used Seurat to process the data with the suggested parameters for this dataset (Butler et al., 2018, 2017). Briefly, genes expressed in 3 or more cells and cells with at least 200 non-zero expression values are retained. After removing outliers (unique gene counts > 2500 and mitochondrial percentage > 5%), we log-normalized the `pbmc3k` data and regressed out technical variations due a number of UMIs and a percentage of mitochondrial gene expression. Among 2638 PBMC samples, we selected 1838 highly variable genes that meet minimum mean and dispersion criteria. K-means clustering is applied on the resulting 2638 PBMC samples containing 1838 genes, using *K* = 8. These 8 clusters contain 346, 290, 177, 16, 186, 33, 1134, and 456 samples with diverse cluster centers (Figure S4).

We directly utilize *K* = 8 clusters of `pbmc3k` data and their corresponding numbers of members to generate an identically sized dataset (2638 PBMC samples with 1838 variable genes) with 10% of noise-only samples. Essentially, we simulated the PBMC dataset such that null samples only containing noise are known. As typical in scRNA-seq data analysis, this PBMC-simulated data is visualized in a t-SNE projection (Figure 4(a)). Without using information about a simulation setup, we applied the jackstraw method to test statistical significance of cluster membership, with *s* = 264 and *B* = 100. P-values, stratified by true and false members of clusters, are visualized in a quantile-quantile plot against a Uniform(0,1) distribution (Figure 4). As theoretically expected, null p-values corresponding to i.i.d. samples follows the dashed diagonal line with a KS p-value of 0.88. In contrast, p-values corresponding to true members are highly significant, deviating strongly downwardly (KS p-value < 2.2*e* − 16).

**Figure 4:**
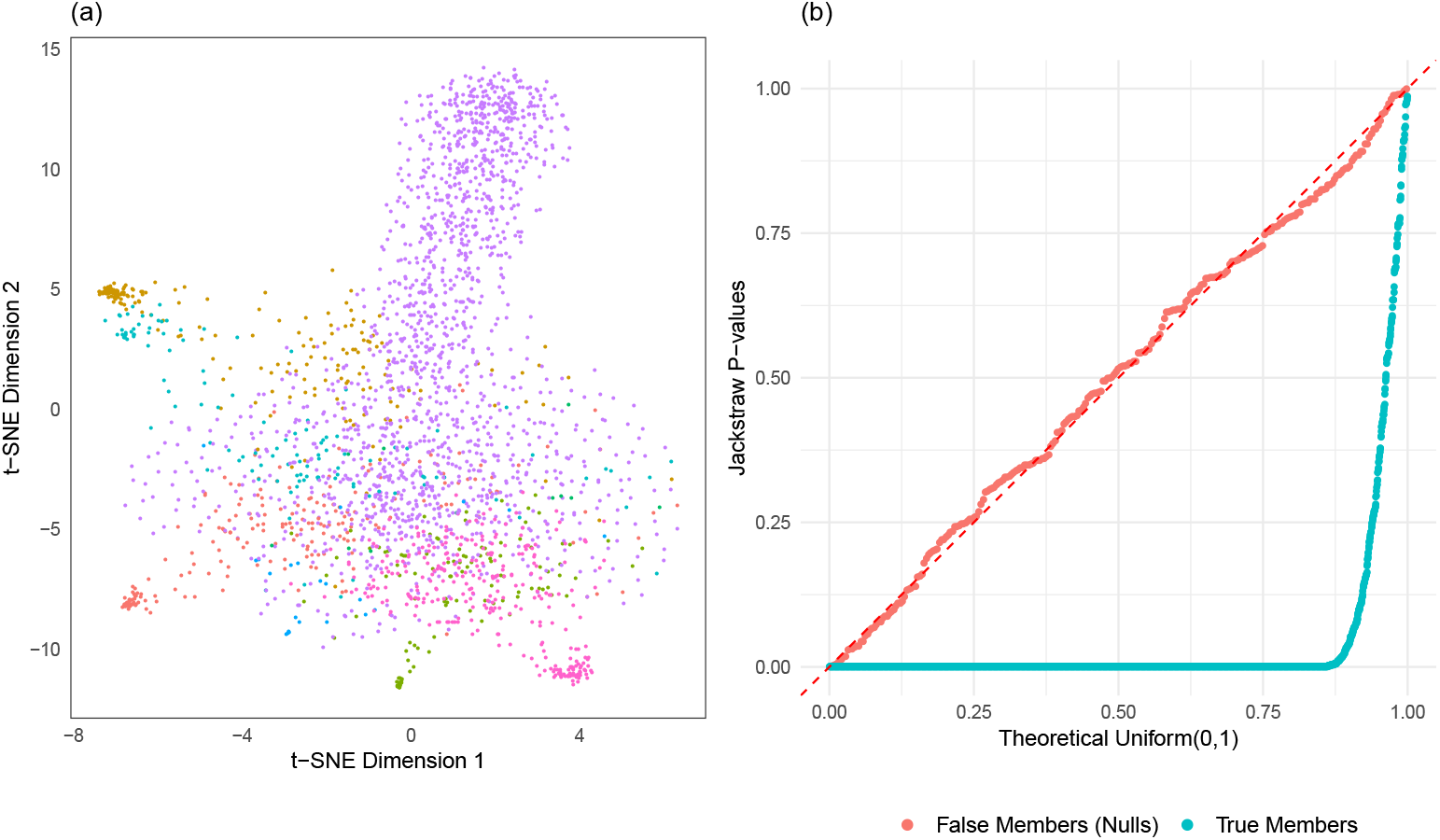
Evaluation of the jackstraw tests using PBMC simulation. A total of 2638 PBMC samples are generated based on ‘pbmc3k’ dataset (10X Genomics) using K=8 clusters where 264 samples are known to only contain noise. *(a)* t-SNE projection displays 8 clusters (different colors) with varying sizes. *(b)* After obtaining p-values from the jackstraw tests for cluster membership, the QQ plot is created stratified by simulated membership status.

### Single Cell Analyses

Whereas conventional microarray and RNA-seq experiments obtain ‘bulk’ gene expression from a sample that contains a large number of cells, scRNA-seq enables more precise and accurate quantification from single cells. Recent studies using high-throughput scRNA-seq measure gene expression from single cell samples, in order to elucidate cellular heterogeneity (Jaitin et al., 2014; Macosko et al., 2015; Zheng et al., 2017). Cell identities are unknown and unlabeled at a single cell level, even though heterogeneous cells are expected to be captured and profiled. As cell identities are computationally determined by clustering, we have developed resampling-based methods to test cell identities. We applied the proposed methods two scRNA-seq datasets from Zheng et al. (2017). Particularly, practical feature selection and visualization techniques are showcased.

### Mixture of Jurkat and 293T Cell Lines

An equal mixture of Jurkat and 293T cell lines (50:50) were sequenced using GemCode by 10X Genomics (Zheng et al., 2017). Jurkat and 293T cell lines are highly distinct, being derived from male and female individuals respectively. While their mixture proportion and cell lines is known, identities of individual single cells that have been captured and profiled by scRNA-seq must be computationally determined. This is a unique experimental data in which *K* = 2 subpopulations indeed exist. Zheng et al. (2017) applied K-means clustering that separates most of samples into *K* = 2 subpopulations. Our proposed methods provide probabilistic measures for how these single cell samples are truly in that subpopulations. Note that the original authors also used known characteristics of two cell lines (such as expression of *CD3D* and *XIST*) to further provide evidence for accurate detection of cellular heterogeneity in a supervised manner.

Following the original analysis pipeline including K-means clustering, we clustered 3381 single cell samples with *K* = 2 based on the top 10 PCs of unique molecular identifier (UMI) counts. The jackstraw tests for those *K* = 2 clusters were conducted with *s* = 100 and *B* = 5000. We found that the jackstraw p-values capture deviation away from two centers, along the 1st PC axis (Figure 5(a)). Using the q-value methodology (Storey and Tibshirani, 2003), the proportion of null samples, which may be assigned to subpopulations by chance, is estimated to be 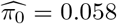. Then, we computed the proposed PIPs from p-values (Figure 5(b)). At PIP < 0.80 (equivalent to 20% local FDRs), 3.4% of 3381 single cells are identified as ambiguous and removed from corresponding clusters. Instead of hard-thresholding the single cell samples at a threshold, we visualized posterior probabilities as levels of transparency in a scatterplot of the top 2 PCs (Figure 5(c)).

**Figure 5:**
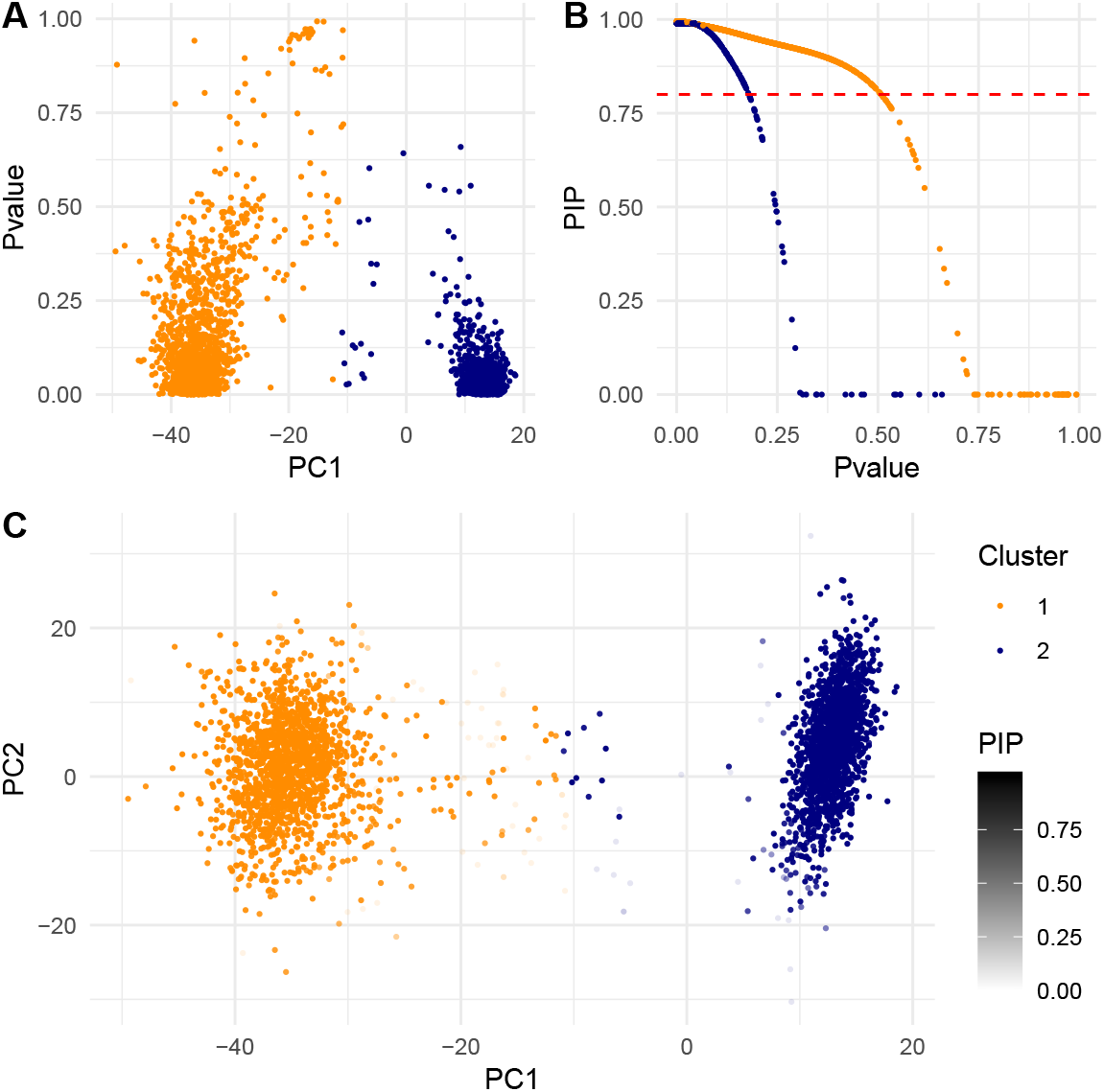
The jackstraw clustering analysis of scRNA-seq data of Jurkat and 293T cells (Zheng et al., 2017). *(a)* Clustering membership p-values are plotted against the 1st PC, which largely separates two cell lines. *(b)* Posterior inclusion probabilities (PIPs) were computed from p-values. At a PIP threshold of 0.80, 3.4% of 3381 single cells would be discharged from corresponding clusters. *(c)* PIPs are visualized as transparency (alpha) levels on the scatterplot of the top 2 PCs. Note that when PIP=0, as appeared in (b), the data point is completely transparent.

Given that a large number of single cells are automatically captured and profiled by a droplet-based platform GemCode, it is possible that a single droplet may contain two or more single cells. Known as doublets or multiplets, they may induce biologically irrelevant gene expression profiles in scRNA-seq studies. Through single nucleotide variant (SNV) detection, Zheng et al. (2017) inferred a 3.1% multiplet rate for this mixture experiment. For ~ 10000 single cells, Zheng et al. (2017) reported > 8% multiplet rates that approximately linearly increase with the recovered cell number. Such contaminations by multiplets are ubiquitous in high-throughput scRNA-seq platforms (Andrews and Hemberg, 2018). The proposed statistical tests effectively identify suspected multiplets with ambiguous cell identities in a completely unsupervised manner. The ambiguous identities of singe cells due to multiplets and other source variations would become increasingly challenging as scRNA-seq becomes more high-throughput, affordable, and widespread.

### Immune Subpopulations among 68K PBMCs

We analyzed scRNA-seq data of 68579 peripheral blood mononuclear cells (PBMCs) that originated from a single healthy donor A (Zheng et al., 2017). Cellular heterogeneity among PBMCs is expected, for which the original authors wanted to characterize immune populations in an unsupervised manner. PBMCs in human are primarily consisted of lymphocytes (T cells, B cells, NK cells), monocytes, and dendritic cells, although several minor subtypes have been investigated. When clustering this 68K PBMC dataset, the proposed statistical tests provide a straightforward evaluation of computationally defined cell identities. P-values and posterior inclusion probabilities (PIPs) are used to remove ambiguous samples and to identify the most relevant samples for immune populations approximated by 10 clusters.

Genes that are expressed in > 1% (69) of observed single cells and single cell samples that have a minimum of 500 genes were retained and processed using Seurat (Satija et al., 2015; Butler et al., 2018). We applied a log-normalization, followed by regressing out technical variations due to batch effects (8 channels), % mitochondrial genes, and numbers of unique molecular identifiers (UMIs). Following the original analysis (Zheng et al., 2017), we selected the 1000 most variable genes by their dispersion among 40507 PBMCs. Then, we obtained the top 50 PCs from applying PCA on variable gene expression profiles of 40507 PBMCs. K-means clustering was applied on the top 50 PCs with *K* =10 which were determined by the sum of squared error (Zheng et al., 2017).

We applied mini batch K-means clustering (Sculley, 2010) on the top 50 PCs obtained from this PBMC data. The mini batch parameters were 10% batch size and 100% of samples for initialization, using K-means++ (Arthur and Vassilvitskii, 2007). The proposed methods for cluster memberships were applied with 10% synthetic null samples and 100 iterations (Figure S5). The proportion of null samples is estimated to be 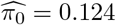. At PIP thresholds of > 0.80 and > 0.90, we found that 34134 (84.2%) and 22407 (55.3%) single cell samples are assigned to their corresponding 10 clusters, respectively. Using an identical perplexity parameter of 30, t-SNE projection after our feature selection improved the original t-SNE projection (Figure 6). Due to a stochastic nature of t-SNE, separate runs may result in different projections. Therefore, we also used the original t-SNE projection using all samples and visualized a subset of samples with high PIPs (Figure S6).

**Figure 6:**
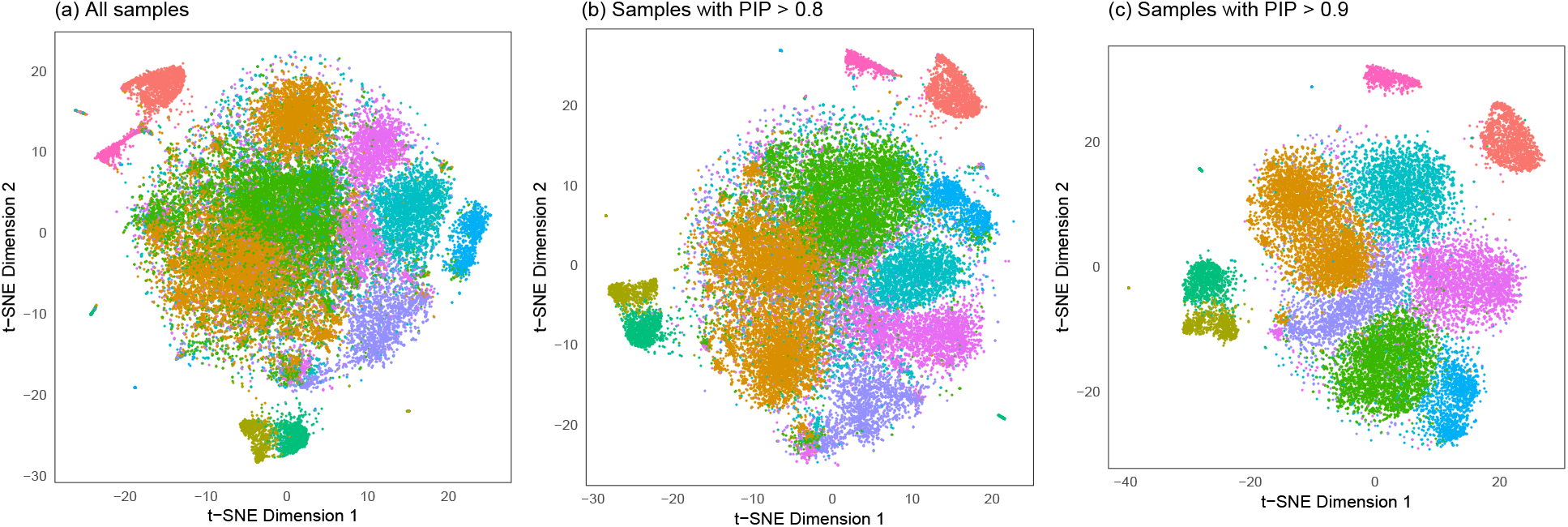
t-SNE projection of PBMCs with proposed feature selections. The top 50 PCs were computed based on the 1000 most variable genes of 40537 samples. Then, we obtained 10 clusters and applied the proposed methods to obtain posterior inclusion probabilities (PIPs) of 40537 samples. *(a)* t-SNE projection based on the top 50 PCs, as per the original analysis (Zheng et al., 2017). After PIPs were computed from the proposed methods, we applied t-SNE on *(b)* 34134 samples with PIPs > 0.8 and *(c)* 22407 samples with PIPs > 0.9. Colors correspond to 10 clusters, which are consistent in three scatterplots.

Note that with 100% of samples for initialization and 10% batch size, mini batch K-means clustering took 3 − 4 seconds for 10 starts and 1000 maximum iterations. K-means clustering on this dataset required 20 − 21 seconds (MacBookPro i5 2.4 GHz). Therefore, we recommend using mini batch K-means clustering and related jackstraw algorithms when a number of single cell samples is over several thousands.

## Discussion

Modern sequencing technologies have rapidly increased the amount and complexity of genomic data. Particularly, recent scRNA-seq platforms enable genome-wide quantification of gene expression in tens of thousands of single cells. Single cells that are captured and profiled may exhibit heterogeneity related to cellular types, spatio-temporal contexts, and environmental stimuli. We are interested in dissecting the cellular heterogeneity by applying unsupervised clustering and assigning cell identities at a single cell level. Our proposed statistical methods evaluate such cluster-based cell identities, by assigning p-values and posterior probabilities of cluster membership to individual single cell samples. The jackstraw approach learns the overfitting characteristics inherent in unsupervised clustering of single cell samples and estimate probabilities that samples truly belong to their assigned subpopulations. We show accuracy and usage of p-values and posterior inclusion probabilities in simulated and scRNA-seq data.

There exists a range of clustering algorithms to automatically assign *m* single cell samples into *K* subpopulations. Given a subset of *m* samples have been assigned to *k*^th^ subpopulation, how can we test if those samples truly belong to *k*^th^ subpopulation? The proposed methods solve this challenge. Our key ingredient is to generate and re-cluster the jackstraw data, in which *s* ≪ *m* synthetic null samples are used to derive the empirical null distribution. Simulation studies based on a latent variable model and a 10X Genomics scRNA-seq data demonstrated accurate operating characteristics that control theoretical error rates. We used rigorous standards, called the joint null criterion, to evaluate statistical significance in multiple hypothesis testing. The proposed statistical tests exhibit a theoretically correct behaviors, that allows one to identify null samples.

For scRNA-seq applications, we considered a mixture of two cell lines and an investigation of immune subpopulations among PBMCs (Zheng et al., 2017). Because of a biological distinction between Jurkat and 293T cell lines, their unlabeled 3381 single cells are well separated in gene expression profiles. Nonetheless, doublets and other ambiguities have been shown to contaminate the data, that become a greater threat to higher throughput studies. The proposed jackstraw tests for cluster membership identify such ambiguous single cell samples without specialized experiments or supervised analysis. Furthermore, we applied the proposed methods on a scRNA-seq data of 68579 PBMCs to examine 10 immune subpopulations (Zheng et al., 2017). Following the original study that demonstrated 10 clusters corresponding to immune subpopulations, we performed feature selection using proposed methods. Emphasizing samples with high PIPs or removing samples with low PIPs are shown to improve visualization of PCA and t-SNE projections.

The proposed methods open new possibilities for selecting canonical cluster members, shrinking cluster centers, and guiding the choice of stable clusters. Our proposed methods enable statistical testing of unsupervised classification at a single cell level, such that the estimated subpopulations can be rigorously used in downstream analyses. The jackstraw test for PCA and related methods (Chung and Storey, 2015) have been used in a variety of genomic studies (Macosko et al., 2015; Satija et al., 2015; Chung et al., 2017; Farré et al., 2015; Jang et al., 2017; Zheng et al., 2017). Complementing this successful approach, we have developed the jackstraw test for clustering. It may be useful to integrate both variants of the jackstraw tests, from selecting highly informative genes to deriving computational cell identities. Because the proposed methods are not limited to genomics, we anticipate its adaptation in other fields of data-intensive science.

### Software Availability

Open-source R package jackstraw is freely available: Stable Release on CRAN https://CRAN.R-project.org/package=jackstraw Development on Github https://github.com/ncchung/jackstraw

## Acknowledgments

This research was supported by the Narodowe Centrum Nauki (National Science Centre) grants 2016/23/D/ST6/03613 and visiting professorship at Prof. Peipei Ping’s lab at the Department of Physiology, UCLA. I would like to thank Prof. John D. Storey of Princeton University for encouragements and discussions.

## Supplementary Materials

**Figure S1:**
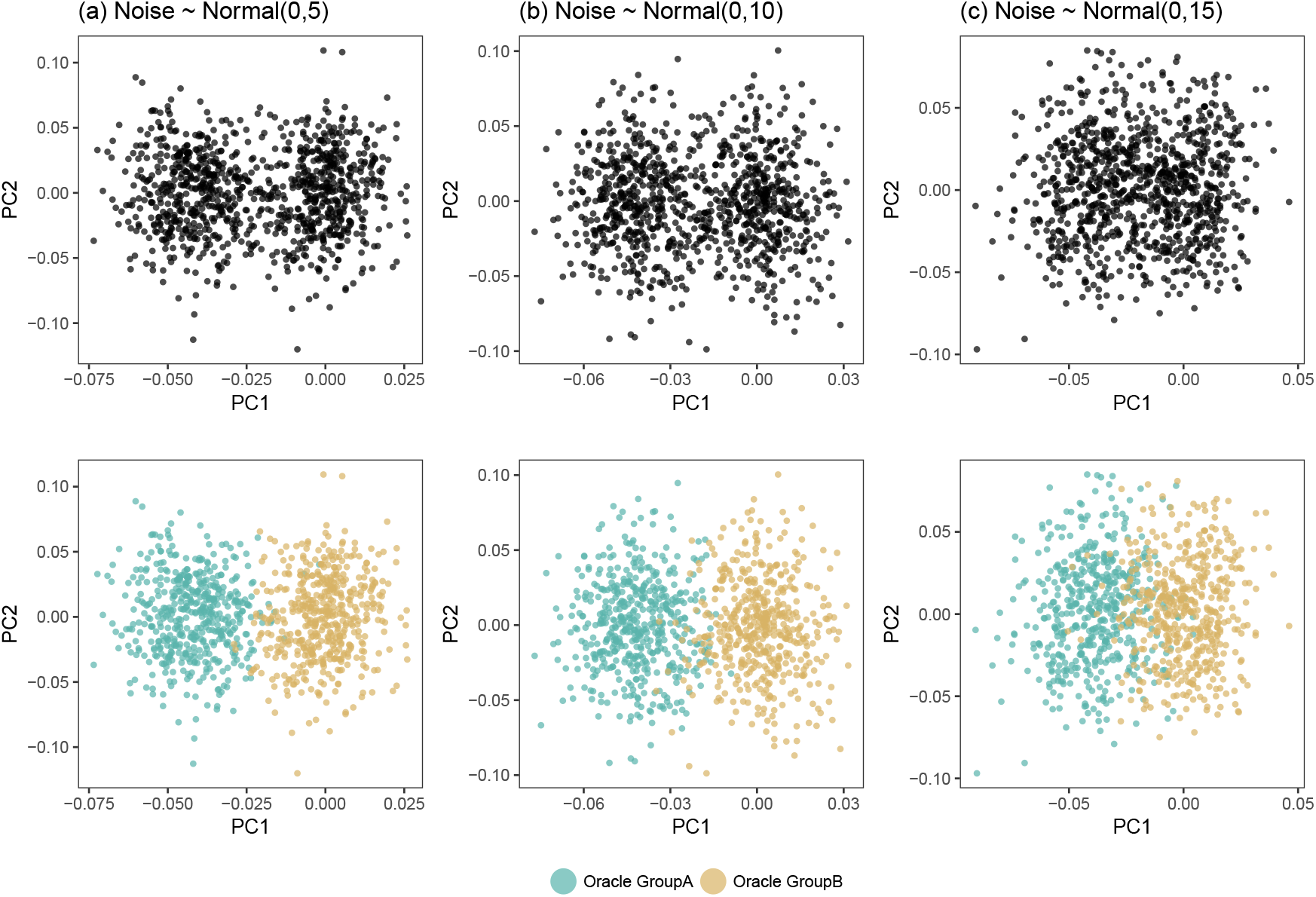
Scatterplots of the top 2 principal components (PCs) from the simulated data. *Oracle Groups* are shown in colors. An increasing level of noise, *σ*^2^ = 5,10,15 brings data samples from two different underlying structures closer together.

**Figure S2:**
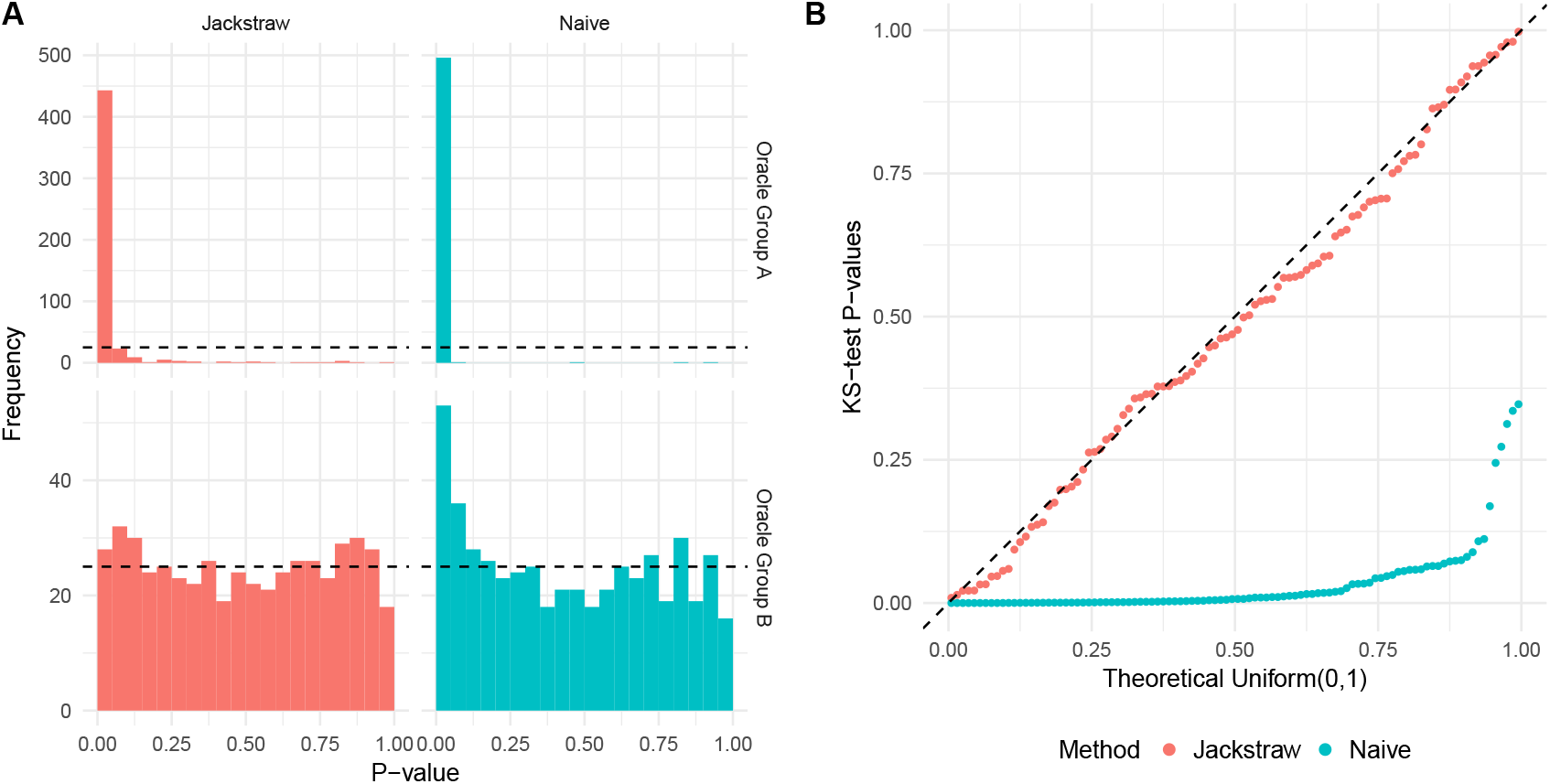
Simulation studies using *σ*^2^ = 5. The jackstraw tests (*s* = 100 and *B* = 5000) or the naive tests are applied without using any information from simulation. *(a)* P-values are shown stratified by *Oracle Groups*, where the naive tests result in an anti-conservative bias. The uniformity of null p-values corresponding to *Oracle Group B* is examined by KS tests, which are independently repeated 100 times. *(b)* The total of 100 independent simulation studies are conducted, and 100 KS-test p-values are plotted against the Uniform(0,1) distribution. The proposed jackstraw tests meet the joint null criterion with a double KS test p-value of 0.81, whereas the naive tests are highly anti-conservative with a double KS test p-value of < 2.2 × 10^−16^.

**Figure S3:**
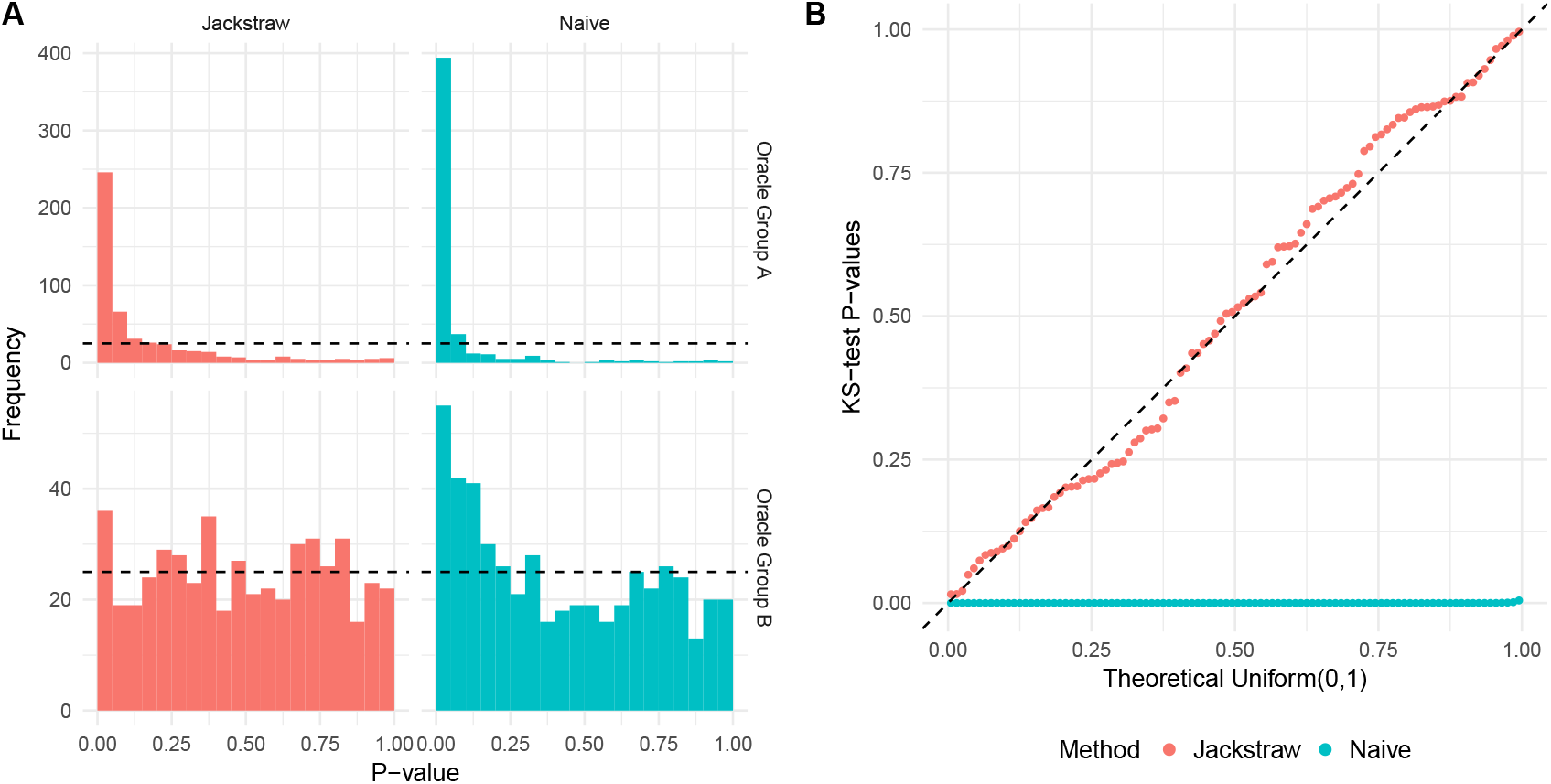
Simulation studies using *σ*^2^ = 15. The jackstraw tests (*s* = 100 and *B* = 5000) or the naive tests are applied without using any information from simulation. *(a)* P-values are shown stratified by *Oracle Groups*, where the naive tests result in an anti-conservative bias. The uniformity of null p-values corresponding to *Oracle Group B* is examined by KS tests, which are independently repeated 100 times. *(b)* The total of 100 independent simulation studies are conducted, and 100 KS-test p-values are plotted against the Uniform(0,1) distribution. The proposed jackstraw tests meet the joint null criterion with a double KS test p-value of 0.67, whereas the naive tests are highly anti-conservative with a double KS test p-value of < 2.2 × 10^−16^.

**Figure S4:**
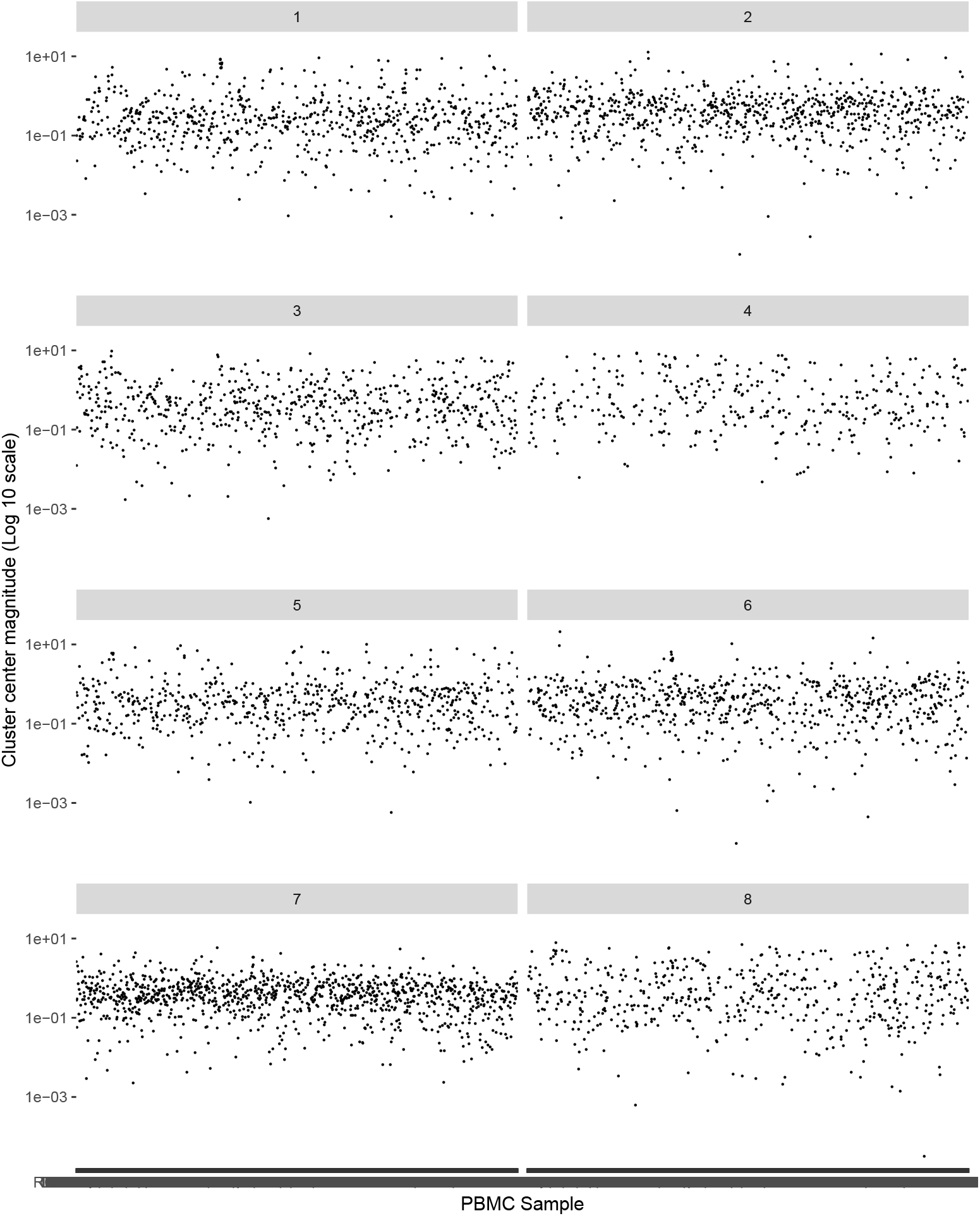
PBMC Cluster Centers. K-means clustering is applied on a gene expression dataset of 2638 PBMC single cells from 10X Genomics’ Chrominum with K=8. These clusters contains 346, 290, 177, 16, 186, 33, 1134, and 456 PBMC samples respectively.

**Figure S5:**
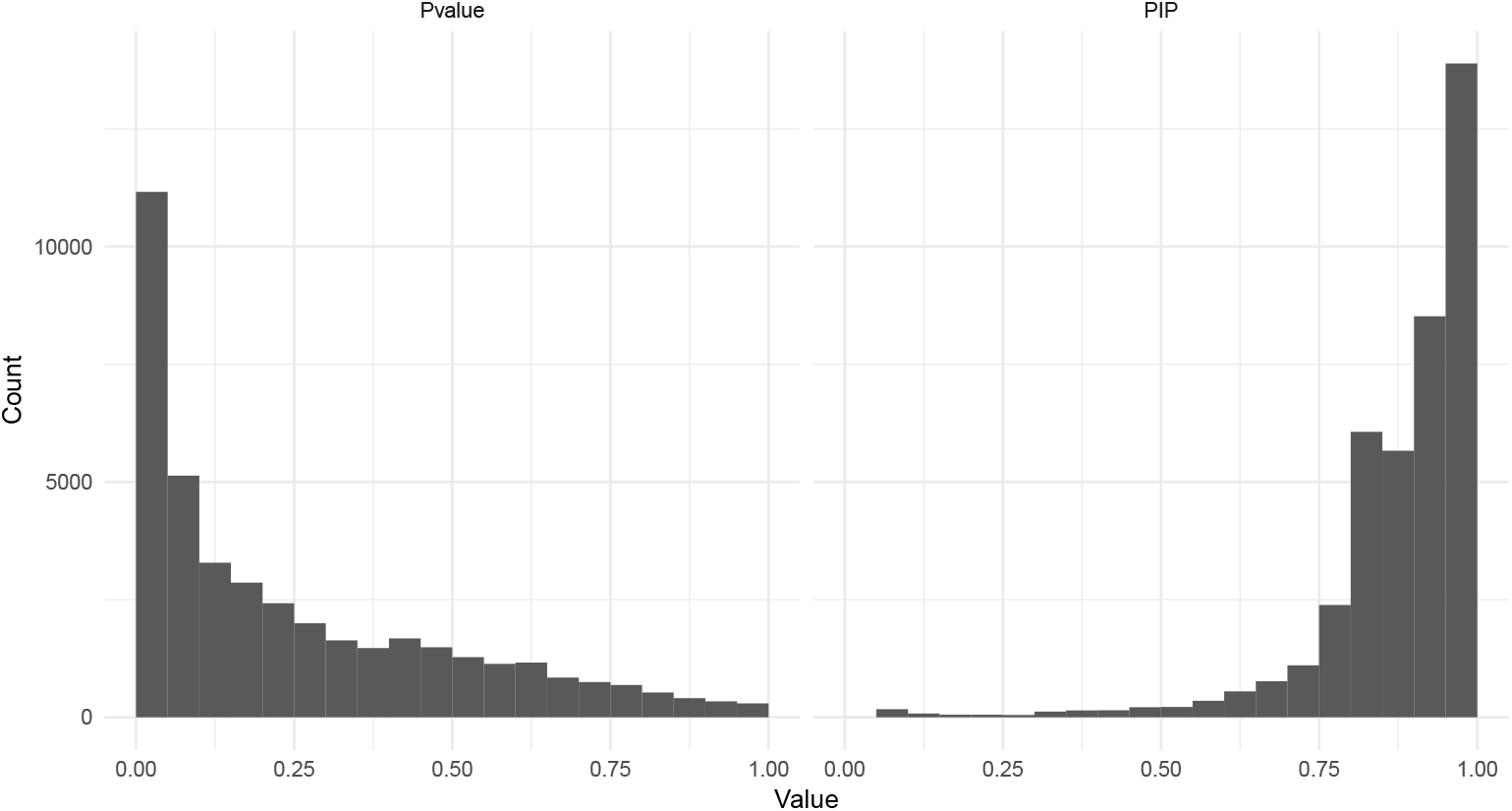
P-value and PIP histograms for applying the proposed methods on 68K PBMC dataset. Mini batch K-means clustering is applied on the top 10 PCs of 1000 variable genes from a processed and normalized 68K PBMC data.

**Figure S6:**
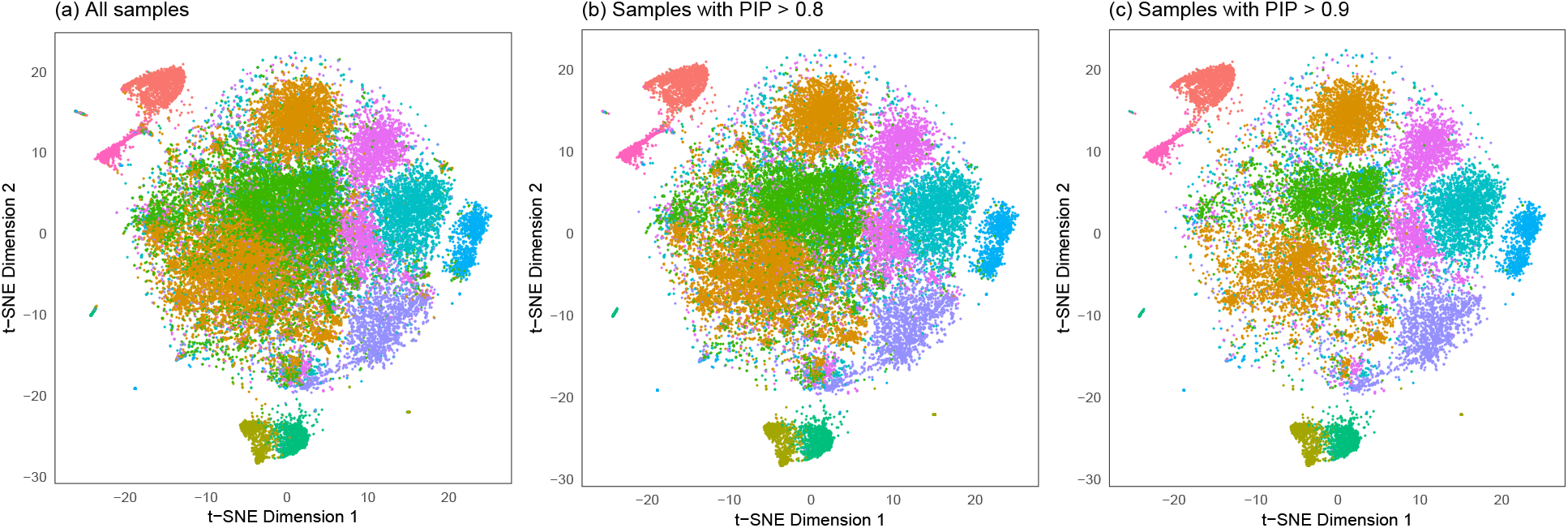
t-SNE projection of PBMCs without recomputing t-SNE. Similar to Figure 6, proposed feature selections for 10 clusters are shown. Instead of recomputing t-SNE, which may stochastically give different projections, the original t-SNE projection using all samples is used in (a)-(c). Then, *(a)* All samples, *(b)* 34134 samples with PIPs > 0.8, and *(c)* 22407 samples with PIPs > 0.9 are shown. Colors correspond to 10 clusters, which are consistent in three scatterplots.

## Footnotes

1 https://support.10xgenomics.com/single-cell-gene-expression/datasets

